# Widespread Historical Contingency in Influenza Viruses

**DOI:** 10.1101/070094

**Authors:** Jean Claude Nshogozabahizi, Jonathan Dench, Stéphane Aris-Brosou

## Abstract

In systems biology and genomics, epistasis characterizes the impact that a substitution at a particular location in a genome can have on a substitution at another location. This phenomenon is often implicated in the evolution of drug resistance or to explain why particular ‘disease-causing’ mutations do not have the same outcome in all individuals. Hence, uncovering these mutations and their locations in a genome is a central question in biology. However, epistasis is notoriously difficult to uncover, especially in fast-evolving organisms. Here, we present a novel statistical approach that replies on a model developed in ecology and that we adapt to analyze genetic data in fast-evolving systems such as the influenza A virus. We validate the approach using a two-pronged strategy: extensive simulations demonstrate a low-to-moderate sensitivity with excellent specificity and precision, while analyses of experimentally-validated data recover known interactions, including in a eukaryotic system. We further evaluate the ability of our approach to detect correlated evolution during antigenic shifts or at the emergence of drug resistance. We show that in all cases, correlated evolution is prevalent in influenza A viruses, involving many pairs of sites linked together in chains, a hallmark of historical contingency. Strikingly, interacting sites are separated by large physical distances, which entails either long-range conformational changes or functional tradeoffs, for which we find support with the emergence of drug resistance. Our work paves a new way for the unbiased detection of epistasis in a wide range of organisms by performing whole-genome scans.

## Introduction

One of the most fundamental questions in biology concerns the emergence of new structures and new functions, in particular at the molecular and genetic level (Lynch, 2007). As such, a large body of experimental work has accumulated over the past decade to unravel the mutational history at the origin of simple phenotypes. For instance, one particular bacterial drug resistance is conferred by five mutations, but out of the 5! = 120 possible ways in which these mutations can accumulate, only a handful of mutational trajectories are evolutionarily accessible (Weinreich *et al*., 2006). This line of work suggests that some mutations are required in order for subsequent mutations to occur. Further work demonstrates that such permissive mutations are not limited to bacteria, as they are also found in vertebrate (Ortlund *et al*., 2007), yeasts (Sorrells *et al*., 2015) and viral systems (Gong *et al*., 2013). However, while such chains of dependent or conditional substitutions —called historical contingency (Mohrig *et al*., 1995; Harms and Thornton, 2014)— are expected to lead to mutational trajectories, their shape and ramifications are not completely elucidated.

As the experimental determination of these trajectories can be tedious, taking > 20 years in the case of the Long-Term Evolution Experiment (Blount *et al*., 2008), computational solutions were sought to reconstruct historical contingencies and the mutational correlations they imply. Initial solutions relied on protein sequence alignments to compute sitewise vectors of amino acid frequencies, from which pairs of co-evolving residues could be identified (Neher, 1994; Taylor and Hatrick, 1994). While this general approach is still used in statistical physics to predict protein folds (Shindyalov *et al*., 1994; Sutto *et al*., 2015), numerous refinements were brought either through the use of metrics such as mutual information (Korber *et al*., 1993; Atchley *et al*., 2000; Gloor *et al*., 2005) or by correcting for shared evolutionary history. One of the first methods to detect correlated evolution while accounting for phylogeny was given in the general context of the evolution of discrete morphological characters (Pagel, 1994). Further leveraging on Schöniger and von Haeseler (1994), the method was quickly extended to analyze RNA molecules by modeling dinucleotides (Muse, 1995; Rzhetsky, 1995) and to map the correlated residues hence detected onto a three-dimensional protein structure (Pollock *et al*., 1999; Poon *et al*., 2007b,a). More recently, a full evolutionary model was proposed to detect epistatic sites in a Bayesian framework (Nasrallah and Huelsenbeck, 2013). However, such approaches rely on complex models and Bayesian computing tools, may scale up poorly with increasingly large data sets (Poon *et al*., 2008; Aris-Brosou and Rodrigue, 2012), and are also generally geared towards detecting positive epistasis (when two mutations increase fitness values).

Here we build on these developments to describe a novel, yet intuitive, statistical method for detecting correlated evolution among pairs of amino acids (AAs) in the typically fast-evolving influenza A virus (Worobey *et al*., 2014). Our method takes inspiration from an approach developed in ecology and aimed at detecting correlated evolution among phenotypic traits (Pagel, 1994; Pagel and Meade, 2006). We validate our approach using a two-pronged procedure based on both extensive simulations and analyses of experimentally-validated data sets (both in viral and in eukaryotic systems). The analysis of several large data sets leads us to reconsider the nature of epistasis in influenza viruses. We find evidence that interacting AAs form networks of sites undergoing substitutions that are most likely to be permissive (*i.e*., contigent), as they occur in a temporal sequence. These networks cover large physical distances among interacting AAs suggesting long-range structural and/or functional effects.

## Materials and Methods

### General approach to detect epistasis

#### Repurposing of BayesTraits

In order to model correlated evolution at the molecular level, we employed the maximum likelihood model implemented in BayesTraits (Pagel, 1994; Pagel and Meade, 2006). See Talavera *et al*. (2015) for a similar model. This model was originally developed as a time-homogeneous Markov process with discrete states in continuous time in order to investigate the coevolution of discrete binary traits on phylogenetic trees. This is achieved by testing whether a dependent model of trait evolution fits the data better than an independent model. The dependent model allows two traits to coevolve as the rate of change at one trait depends on the state at the other trait, while the independent model does not place any restriction on rates of change (Pagel, 1994) (see Figure S1). In both cases, the likelihood function is optimized by summing (integrating) over all unobserved pairs of character states at internal nodes. As these two models are nested, a likelihood ratio test can be employed for model selection. The test statistic, twice the log-likelihood difference, is assumed to follow a *χ*^2^ distribution with four degrees of freedom under the null hypothesis (independence).

This general framework further assumes a phylogenetic tree, with a known topology and branch lengths proportional to the amount of evolution separating each node. Both assumptions can be addressed as described below, either by a bootstrap analysis, or by resorting to trees sampled from their posterior distribution.

#### Data recoding

The model described above was implemented for binary traits. Here however, our goal is to analyze the coevolution of pairs of sites (DNA or AA) along a sequence alignment. In the case of proteins, each site has twenty possible AAs, which represent the states of our system. To avoid resorting to tensor kernels, data recoding is therefore necessary to reconcile the data with the approach. In this context, we evaluated two strategies. First, AAs were partitioned according to their physicochemical properties. Binary properties (side-chain group types) were naturally recoded “0” / “1”. For those with *k* > 2 states, we compared each of the *k*_i_ states against the other ones (*k*_*j*_ where *i* ≠ *j*). For instance, in the case of charge, we first assigned state “0” to negative and state “1” to non-negative AAs, and circled through the two other states (Figure S2). This recoding was based on the physicochemical properties as implemented in the R package `protr` ver. 0.2-1 (Xiao *et al*., 2014). However, results from this first method failed to find any significantly correlated pairs of mutations when applied to our first dataset. Hence a second recoding method was devised and used thereafter.

In our second recoding method, AAs were classified as either being in the outgroup or ingroup consensus state: at each position of an alignment, the outgroup state was defined as the majority-rule consensus AA present in a clade used to root the tree. Here, this rooting clade was defined either as the one containing the oldest sequences and this clade was removed for downstream analysis, or as the basal clade in a relaxed molecular clock analysis (see below). The ingroup state was then defined as any AA that differs from the consensus AA in the outgroup. Within each column of the alignment, sites sharing the outgroup state were all recoded as “0”, while those sharing the ingroup state were recoded as “1”.

#### Code optimization

To discover pairs of AAs that are potentially interacting, we need to run the above model on all 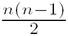pairs of sites in an alignment of length *n*. This computation can be prohibitively long even with an alignment of modest size. For instance, the influenza H3N2 nucleoprotein has 498 AAs, which leads to analyzing 123,753 pairs of sites under both the dependent and the independent model (*i.e*., 247,506 models need to be run). To decrease the computational cost, we first compressed the alignment into site patterns (Yang, 2006) (p. 105). Then, as pairs of sites can be independently compared, we parallelized the code using R’s `foreach` ver. 1.4.2 and `doMC` (ver. 1.3.3) (Analytics and Weston, 2013) packages to take advantage of multicore / multiprocessor architectures (Figure S3). To account for multiple testing, we computed the False Discovery Rate (FDR) according to the Benjamini-Hochberg (BH) procedure (Benjamini and Hochberg, 1995). We used R (v3.0.2) for all analyses (Team *et al*., 2013). In the analyses presented below, the pairs of AAs identified to be interacting were subsequently mapped on three-dimensional protein models predicted by homology modeling with SWISS-MODEL (Biasini *et al*., 2014), and plotted using KiNG (Chen *et al*., 2009).

### Validation based on simulations

To validate our method for detecting correlated pairs of sites, we followed two approaches: an extensive simulation study and the analysis of data sets in which epistasis was experimentally confirmed. We first present our simulation strategies that, in order to avoid biasing our results, were based on two different frameworks of simulating correlated evolution. The Supplementary Text presents additional simulation results.

#### Simulations with PHASE

First, sequences were simulated with PHASE 2.0 (Gowri-Shankar and Jow, 2006), which allowed us to simulate two categories of sites: those that evolve independently and those that evolve in a correlated manner. Sites that evolve independently (main sequence of length *l*_*i*_) were simulated using the General Time-Reversible (GTR) model of nucleotide substitution (Tavaré, 1986) (see also Aris-Brosou and Rodrigue, 2012). Both nucleotide frequencies and transition probabilities were arbitrarily set to reflect published values for Pseudomonas aeruginosa (Garrity *et al*., 2004), although this particular setting has no impact on downstream analyses. With PHASE, sites that evolve in a correlated manner (epistatic sites *l*_*e*_) were simulated under the RNA7D model of dinucleotide substitution (Tillier and Collins, 1998). We modified the transition probabilities such that double substitutions occurred for 95% of changes, single substitutions for 5%, while mismatch substitutions were not permitted. In the results presented below, equilibrium dinucleotide frequencies were all assumed to be equal (set to 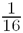). Because influenza A viruses typically exhibit dinucleotide bias (Greenbaum *et al*., 2008), which could lead to false positives, we also performed simulations under the RNA16 model with a dinucleotide bias matching that of influenza A viruses (see Supplementary Text). As total sequence length was determined ahead of time (part of our factorial design), when simulating correlated evolution, we first simulated a number *l*_*e*_ of pairs of epistatic sites, which were then concatenated to the nucleotide sites simulated under the independent model of length *l*_*i*_ to obtain a final sequence of length *l*_*s*_ = 2*l*_*e*_ + *l*_*i*_. Where correlated evolution was not simulated *l*_*s*_ = *l*_*i*_.

#### Simulations with Coev

Our second simulation strategy employed Coev (Dib *et al*., 2014), which produces correlated evolution by randomly defining a dinucleotide profile (two-letter nucleotide states, *e.g*.: AA, AT, CG, *etc.*) and simulating evolution under the Jukes-Cantor substitution model (Jukes and Cantor, 1969). The transition rates for each nucleotide of the dinucleotide may be set independently; however, as we have no a priori reason to assume either evolves with a different rate, we left these rates equal to their default setting. The strength of selection for the defined profile is governed by the ratio of the parameters *d* and *s* (*i.e*., *d/s*), which represent the likelihood of a dinucleotide evolving toward (*d/s* > 1) or away (*d/s* < 1) from the defined dinucleotide profile. When *d/s* = 1, there is no selection and sites evolve independently; we varied the strength of selection from independent (*d/s* = 1) to the default setting offered by Coev-web (Dib *et al*., 2015) (*d/s* = 100), with intermediate strengths (*d/s* = {2, 33, 66}). PHASE was still used to simulate sites that evolved independently (*l*_*i*_) to be consistent in our use of a GTR substitution model.

#### Sensitivity and specificity

In both sets of simulations, we assessed the impact of branch lengths, tree shape, and number of sequences, both in the presence and absence of epistasis (Figure S4B). With PHASE, branch lengths (b) were varied across a *log*_2_ scale for *b* ∈ (– 12, –1). To reduce possible factorial combinations, all branches of a single simulated tree were the same length. Two tree shapes were used, simulated tree topologies (*τ*) being either symmetrically bifurcating or pectinate (Figure S4). All simulated trees were ultrametric. Each tree contained a number of sequences (*n*_*s*_) equal to 16, 32, 64 or 128. This led to a full factorial design containing 192 simulation conditions (12*b* × 2*τ* × 4*n*_*s*_ × 2 for with or without epistasis). Each simulation condition was replicated 100 times and *l*_*s*_ was set to 100 bp. When epistasis was simulated, the number of epistatic pairs was set to 3. An ANOVA was used to assess the significance of each of these factors. All tests were conducted at the *α* = 0.01 (1%) significance threshold.

With Coev, branch lengths (*b*) were varied across a *log*_2_ scale for *b* ∈ {– 12, –10, –8, –6 – 4}. Each tree contained a number of sequences (*n*_*s*_) equal to 32, or 128. The same two tree shapes (*τ*) were used, being either symmetrically bifurcating or pectinate. This led to a full factorial design containing 100 simulation conditions (5*b* × 2*τ* × 2*n*_*s*_ × 5*d/s*). Each simulation condition was replicated 100 times and *l*_*s*_ was set to 100 bp. When epistasis was simulated, the number of epistatic pairs was set to 3, as under PHASE. All tests were here again conducted at the *α* = 0.01 (1%) significance threshold unless otherwise stated.

### Validations based on previous evidence

#### Previous computational analyses

As a first validation of our approach on actual data, we reanalyzed those studied in a previous computational study (Kryazhimskiy *et al*., 2011). The four data sets obtained from S. Kryazhimskiy (*pers. comm*.) consist of: 1,219 HA and 1,836 NA sequences from H1N1 viruses, as well as 2,149 HA and 2,339 NA sequences from H3N2 viruses. These data sets are here denoted KDBP11-H1 (HA in H1N1), KDBP11-N1 (NA in H1N1), KDBP11-H3 (HA in H3N2) and KDBP11-N2 (NA in H3N2), respectively.

#### Experimental evidence

As computational studies make predictions that are not always tested or validated, we reanalyzed data sets in which epistasis was experimentally confirmed. A number of recent studies reported evidence for epistasis and we present results on three of these.

First, we reanalyzed data published by Gong and collaborators (Gong *et al*., 2013) which comprised 424 H3N2 human influenza A nucleoprotein (NP) sequences spanning 42 years between 1968-2010. This gene was originally chosen because it evolves relatively slowly and is hence amenable to experimental validation as all substitutions can be easily tested by site-directed mutagenesis (Gong *et al*., 2013). The viral sequences that they used were downloaded from the IVR database (Bao *et al*., 2008). This data set is here denoted Gong13NP.

We also retrieved a second data set as analyzed by Duan and collaborators (Duan *et al*., 2014), who used an alignment that contained 1366 human influenza A H1N1 neuraminidase (NA) collected between 1999 and 2009; as in their analysis, the 2009 H1N1 pandemic sequences were excluded. This data set is denoted Duan14NA.

Finally, a recent study performed a combinatorial analysis based on mutations of the eukaryotic tRNA^*ArgCCT*^ gene to detect epistasis (Li *et al*., 2016). We ran our computational analysis on an alignment of the eukaryotic tRNA gene constructed in a manner similar to Li and collaborators (SI text: “Phylogenetic data of the tRNA genes”): we downloaded all available eukaryotic tRNA^*ArgCCT*^ genes from GtRNAdb (Chan and Lowe, 2016), aligned the sequences with the cmalign tool from Infernal (Nawrocki and Eddy, 2013), and kept the 75 nucleotide region that spanned the conserved portion within Saccharomyces sp. As our method requires a phylogenetic tree, we used a phylogeny of eukaryotes(Hedges *et al*., 2015), keeping only those tips corresponding to the taxa in our alignment. Lastly, as the branch lengths of this tree were in millions of years, we rescaled these to expected number of substitutions using the best available eukaryotic tRNA molecular clock (Soares *et al*., 2009). While Li *et al*. (2016) did not perform a fully exhaustive analysis of all pairs of sites, and our analysis requires sites to be polymorphic, we present our comparison for those pairs of sites tested in both of the analyses. For this analysis only, we used a statistical threshold of *α* = 0.05 as in Li *et al*. (2016). This data set is denoted Li2016.

### Nature of correlations in influenza evolution

After performing these validations, both on simulations and on experimentally-validated data, we set out to investigate the nature of these interactions by testing two hypotheses. First, we revisited the work by Koel and collaborators (Koel *et al*., 2013) that experimentally validated the existence of AA substitutions involved in changes of antigenic clusters. For this, we used 877 H3N2 human influenza hemagglutinin (HA) sequences as in Koel *et al*. (2013), collected between 1968 and 2003, to test if these substitutions responsible for antigenic changes also showed evidence for correlated evolution. This data set is denoted Koel13HA.

Second, we tested if our statistical approach could detect some evidence for correlated evolution at pairs of sites involving the S31N substitution, which is responsible for conferring resistance to the anti-influenza drug Adamantane (Abed *et al*., 2005). For this, we retrieved 668 H3N2 human influenza A matrix protein 2 (M2) sequences that were collected between 1968 and 2003. This data set is denoted Adam03M2.

### Phylogenetic analyses

Maximum likelihood was employed to estimate phylogenetic tree for the KDBP11-H1, KDBP11-N1, KDBP11-H3, KDBP11-N2, Gong13NP, Duan14NA and Adam03M2, alignments with FastTree ver. 2.1.7 (Price *et al*., 2010) under the WAG +Γ_4_ model to account for among-site rate variation. Trees were rooted using the earliest sequences, which were AAD17229/USA/1918, CAA24269/Japan/1968, AAF77036/USA/1918, ABI92283/Australia/1968, Aichi68-NPA/Aichi/2/1968, A/Victoria/JY2/1968 and A/Albany/17/1968, respectively. The R package `APE` was used to visualize the trees (Paradis, 2006). For the Koel13HA data set, a rooted tree was reconstructed using BEAST ver. 1.8.0 under a relaxed molecular clock assuming an uncorrelated lognormal prior (Drummond *et al*., 2006) and a constant-size coalescent prior under the FLU +Γ_4_ substitution model. Analyses were run in duplicate to check convergence, for a total of 100 million steps with a thinning of 5,000; log files were combined with LogCombiner after conservatively removing the first 10% of each chain as a burn-in period, as checked with Tracer ver. 1.5 (tree.bio.ed.ac.uk/software/tracer).

Phylogenetic uncertainty was taken into account by running our algorithm on bootstrapped trees (or trees sampled from the posterior distribution). Because only the Gong13NP data set showed a large proportion of SH-like aLRT (Anisimova and Gascuel, 2006) node support values in the low (0.0, 0.8) range (Figure S5-S12), the results of the bootstrap analyses are only shown for this case. All data sets and the AEGIS script (Analysis of Epistasis & Genomic Interacting Sites) used in this work are available at github.com/sarisbro/AEGIS.

## Results and Discussion

### Simulation studies

#### Excellent specificity but mediocre sensitivity

As a first means to validating our approach, we conducted a fully factorial simulation study. In order to avoid biasing our results, the simulation models differ from the analysis models. Our simulation results with PHASE demonstrate that alignments containing fewer than 32 sequences have a poor ability to detect epistasis, with a sensitivity generally ≤ 20% (Figure S13). For this reason, we henceforth focus on alignments of at least 32 sequences.

The specificity (*Sp*) of our approach is never below 99.9999%, which occurs for the symmetric tree shape with the largest number of sequences and rather short branch length of 2^−6^ substitutions per site (Figure 1A). Variation in specificity was so low that we report the value on a – *log*_10_(1 – *Sp*) scale in order to emphasize the performance of our approach against thresholds set by the minimum, mean and maximum number of pairwise comparisons among replicates. These thresholds represent the number of true negatives calculable in our analyses (*i.e*., (# pairwise comparisons – # epistatic pairs)/# pairwise comparisons) and thus performance above these thresholds demonstrates that high specificity results from low false positive detection rate, and not simply from a large dataset with unduly large numbers of true negatives. In spite of the low variation in specificity, the number of sequences (*n*_*s*_) explains most of the variance (Table S1). While we find a two-way interaction with tree shape (*τ*) and branch length, there was no direct interaction between *n*_*s*_ and either branch length or *τ* (Table S1). We find a three way interaction between branch length, *τ*, and *n*_*s*_, which reinforces the idea that while the number of sequences dominates the specificity of our method, the amount of evolutionary time (sum of branch lengths over topology) remains an important factor (see Figure 1A).

**Figure 1.**
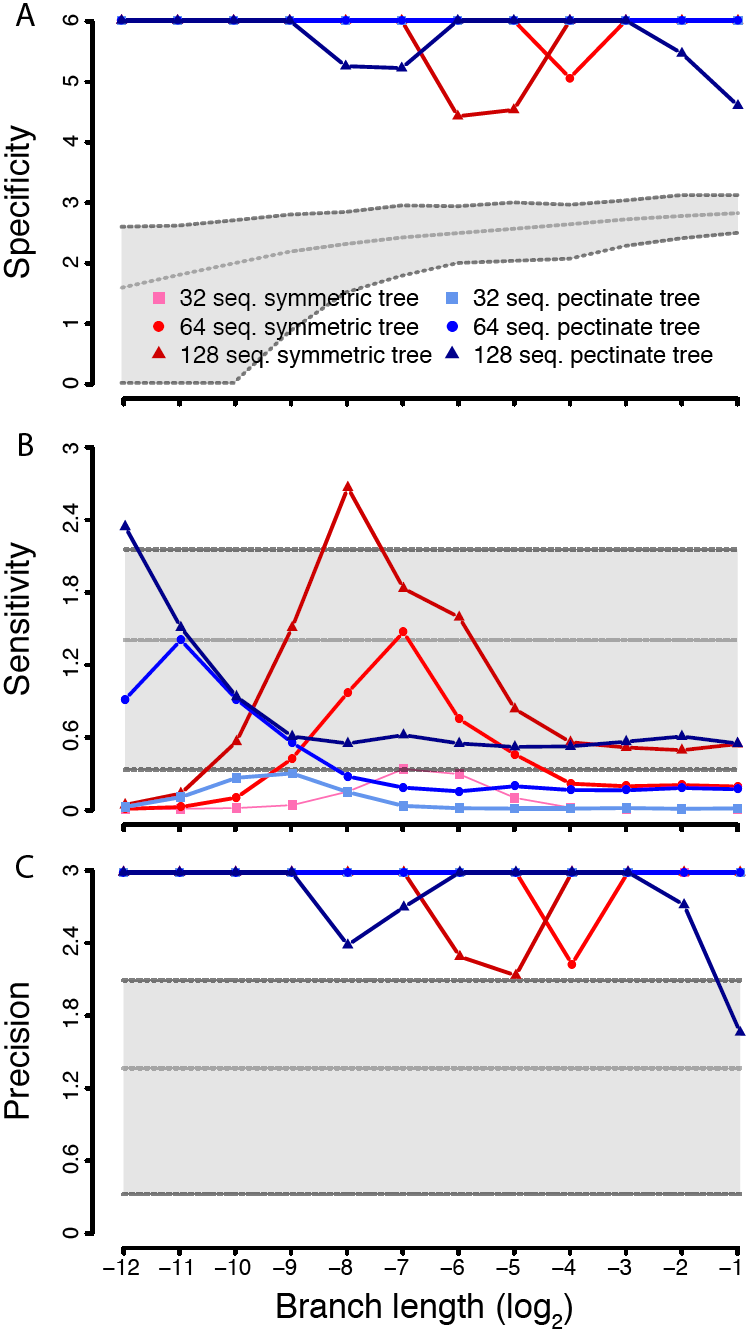
Specificity, sensitivity and precision results, from simulated data, of our novel epistasis detection method. Results are shown for alignments with 32 (squares), 64 (circles) and 128 (triangles) sequences. Tree shapes are color-coded (symmetric in red; pectinate in blue). Branch lengths were varied on a log_2_ scale. All y-axes show the mean of –*1og*_10_(1 – *summary statistic*) to highlight performance of our method as a *summary statistic* approaches 1; when this value is 1, we arbitrarily assigned it a value 10% larger than the largest finite value within the set. Panel (A) shows specificity (true negative rate) with a grey shaded polygon which illustrates thresholds for excellent specificity in parameter space. The thresholds were established by subtracting three (the number of epistatic pairs simulated) from the minimum (lower dashed line), mean (middle dashed line) and maximum (upper dashed line) number of pairwise comparisons performed across all simulations with the same branch length. These thresholds represent the number of calculable true negatives and allow us to demonstrate our method’s excellent specificity. Panels (B) and (C) show sensitivity (true positive rate) and precision (positive predictive value) respectively. Each panel includes a grey shaded polygon which illustrates thresholds for 50% (lower dashed line), 95% (middle dashed line) and 99% (upper dashed line) detection. These thresholds were arbitrarily chosen to demonstrate idealized benchmarks of performance.

While specificity is excellent, we find that sensitivity is mediocre (Figure 1B), which is consistent with previous studies (Poon *et al*., 2007b). Our approach’s sensitivity is a function of branch lengths and the number of sequences (Table S1), with the existence of an optimal branch length where sensitivity could become excellent, reaching close to 100%, before degrading quickly again (on a log scale). This optimal response reflects that short branch lengths carry no information while long ones have random site patterns and thus convey no information (Yang, 1998). Pectinate trees show an optimum for shorter branch lengths than symmetric trees, and an ANOVA confirms an interaction between tree shape and branch length with respect to sensitivity (Table S1). Lastly, we note that larger alignments appear to have better sensitivity (Figure 1A) and that this does have an interaction with tree shape. We note that there exists large variance in sensitivity, due to our method of simulating epistasis, but have omitted error bars from Figure 1 for reasons of clarity. Due to the low sensitivity of our method, we further investigated the prevalence of false positives by plotting precision (Figure 1C). From this figure we find that less than 1% of positives are false except when branches are long.

Because dinucleotide bias, as typically found in influenza A viruses, could lead to false positives, we also assessed the impact of this parameter on the performance of our approach. We mimicked bias as found in the four KDBP11 data sets, and employed the full RNA16 model in PHASE to perform simulations similar to those under GTR as above. Our results show that both specificity and precision are unchanged, while sensitivity appears to be decreased. As we did not change the method simulating dependent sites, we did not expect sensitivity to be affected. After thorough review of our data we can only conclude that his difference results from the large variance in sensitivity resulting from the method by which epistasis is simulated. Altogether, dinucleotide bias does not lead to an increase in false positives (Figure S14).

To further confirm that our simulation results are not biased by our method of simulating epistasis, we compared our detection method to simulated evolution generated by Coev (Dib *et al*., 2014). These results (see Supplementary Text) show that both specificity and precision are comparable with the PHASE simulations, while sensitivity is reduced (Figure S15). These findings reflect the particularities of how epistasis was simulated and confirm the excellent specificity and precision of our method. Importantly, these properties are found even when epistasis is simulated at very weak levels (Figure S15).

Note that unlike the real data sets analyzed in the next subsection, all these simulations are based on a state space of size four (DNA) and not 20 (AAs): the effect of this reduction of the state space in the simulations is to lead to fewer unique site patterns, and hence larger proportions of false positives in DNA data than in AA data. Our DNA-based simulations show however that false positives are already well-controlled.

### Real data analyses

#### Limited overlap with previous computational results

As a first evaluation of our algorithm on real data, we reanalyzed four large data sets (> 2000 sequences) previously analyzed with another computational method designed to detect epistasis (Kryazhimskiy *et al*., 2011). Briefly, this method consists in mapping mutations on the phylogeny by parsimony, in order to then infer pairs of sites that are candidates for correlated evolution, before determining if mutations at each pair are temporally “close enough” to be considered as interacting. Unlike our approach, no data recoding is necessary; conversely, no mapping is involved in our method as the likelihood function describing changes of states at pairs of sites plays this role.

Here, the AA data were recoded as outgroup / ingroup character states to match the above simulations: at each position of the alignment, the “outgroup” state represents the majority-rule consensus AA in the outgroup sequences and the “ingroup” state represents all the AAs types that differ from the outgroup consensus AA. Overall, our approach not only detects fewer epistatic sites than the previous method, but the two approaches also show limited overlap (Figure S16). This marginal overlap is even found before FDR correction (Figure S17, Tables S3-S6), so that lack of power is an unlikely explanation of the difference. Our extensive simulations suggest that this difference may be the result of the low-to-average sensitivity and of the excellent specificity / precision of our method, so that the AAs pairs that we detect may actually be coevolving sites among the truly epistatic pairs. This result begs the question as to whether the few pairs of sites we detect would have any experimental evidence supporting epistasis.

#### Detected pairs are almost all experimentally-validated

Because the previous data sets were only examined from a computational point of view, we turned to additional data that have been experimentally validated.

First, we analyzed the Gong13NP data set of 424 influenza NP protein sequences (Gong *et al*., 2013). As this is the smallest data set in our study, we assessed two ways of recoding the data into binary character states. We first partitioned AAs according to their physicochemical properties. With this recoding strategy, four pairs of sites were detected before FDR correction (Table S7), but none of them matching the pairs detected by Gong and collaborators. Note that after FDR, no interactions were significant (Table S7).

The estimated phylogenetic tree for this data set shows a pectinate-like (asymmetric) shape and short branch lengths (Figure 2) – even if H3N2 viruses exhibit a more punctuated mode of evolution than H1N1 viruses (Sandie and Aris-Brosou, 2014). Our simulation results show that, under these conditions, with > 100 sequences, we can expect a sensitivity ≥ 80% (Figure 1A), so that the Gong13NP data set fulfills all the conditions for detecting true interactions. This suggests that even if the physicochemical recoding seemed *a priori* to be a good idea, capitalizing on the chemistry of life, it wastes statistical power on multiple three-way tests (Figure S2).

**Figure 2.**
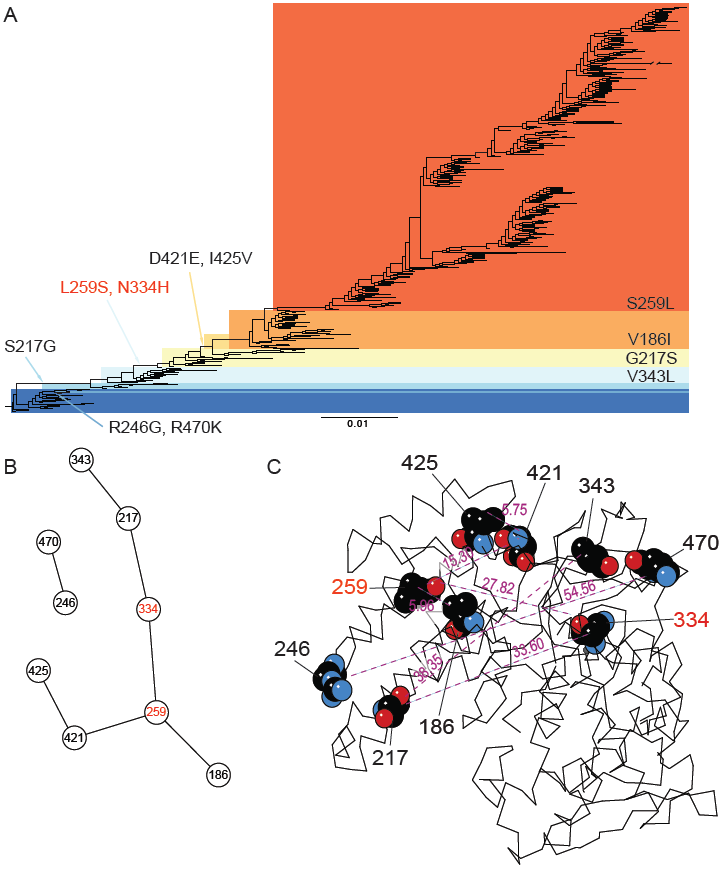
Epistatic pairs of amino acids detected in the Gong13NP data set with the outgroup / ingroup recoding. (A) The epistatic mutations that we detected are plotted on the NP phylogenetic tree. The substitutions in red were experimentally validated (Gong *et al.*, 2013). (B) Chained epistasis of interacting AAs. (C) The epistatic sites are mapped on three dimensional NP protein structure (based on template 3ZDP). The numbers show the AA positions experimentally validated (in red) and those detected only in this study (in black). The numbers in purple show the physical distance between epistatic sites (in Å).

To better mimic our simulation conditions, we then recoded AAs as outgroup / ingroup states. With this recoding strategy, seven pairs of sites were detected: 259:334, 421:425, 246:470, 217:334, 217:343, 259:421 and 186:259 (Figure 2A, Table S8). Gong et al. (2013) detected 259:334 as their strongest signal, as well two additional pairs (384:65 and 280:312), with a weaker signal. However, we detected six additional pairs of sites that were never shown to be epistatic. These could either be false positives —but again our simulations suggest that our approach is extremely specific (Figure 1B)— or simply coevolving pairs of sites that are not epistatic (coevolution is necessary but not sufficient for epistasis to exist), or that are evolving under negative epistasis.

While we find multiple pairs of correlated sites, almost all these pairs are linked to the experimentally confirmed L259S and N334H substitutions (Figure 2B), hereby forming a network of interacting sites. Can we say anything about the nature of these interactions? If they were physical, we would expect that interacting sites would be in close spatial proximity on the folded protein, as in the case of compensatory mutations in RNA molecules (Kimura, 1985; Chen *et al*., 1999). However, the spatial distribution of these epistatic pairs of sites on the protein structure does not conform to this prediction: while two residues are considered physically linked when their distance is ≤ 8.5 Å (Atilgan *et al*., 2004), we find that the average distance between interacting AAs is 25.9 A (*sd* = 18.1; Figure 2C). This distribution strongly suggests that chained epistasis is not linked by spatially-close physical interactions (Figure S18), so that thermodynamic (Thomas *et al*., 2010) or compensatory changes of the 3D structure (Weinreich *et al*., 2006) act at very long spatial ranges.

To further validate our approach, we also analyzed the Duan14NA data set, comprising 1366 NA sequences of pre-pandemic H1N1 viruses (Duan *et al*., 2014). We identify the epistatic pair 275:354 (Figure 3, Table S9), which was experimentally proven to confer oseltamivir resistance and that dominated the population in 2008-2009 (Duan *et al*., 2014). In their study, Duan and collaborators showed that D354G was the main mutation responsible for maintaining the function of NA after alteration of enzyme activity by H275Y (H274Y in N2 numbering), so that this is potentially the strongest existing interaction. However, our approach fails to identify the five other mutations (V234M, R222Q, K329E, D344N and D354G) that were further identified by Duan and collaborators to be interacting with H275Y. The estimated H1N1 tree is more symmetrical in shape than the one estimated for the Gong13NP data and has shorter average branch lengths (see scale bar in Figure 3). Our simulation results suggest that in this case, the sensitivity of our approach can be very small (Figure 1A). This low sensitivity might explain why we fail to detect the five additional sites interacting with position 275.

**Figure 3.**
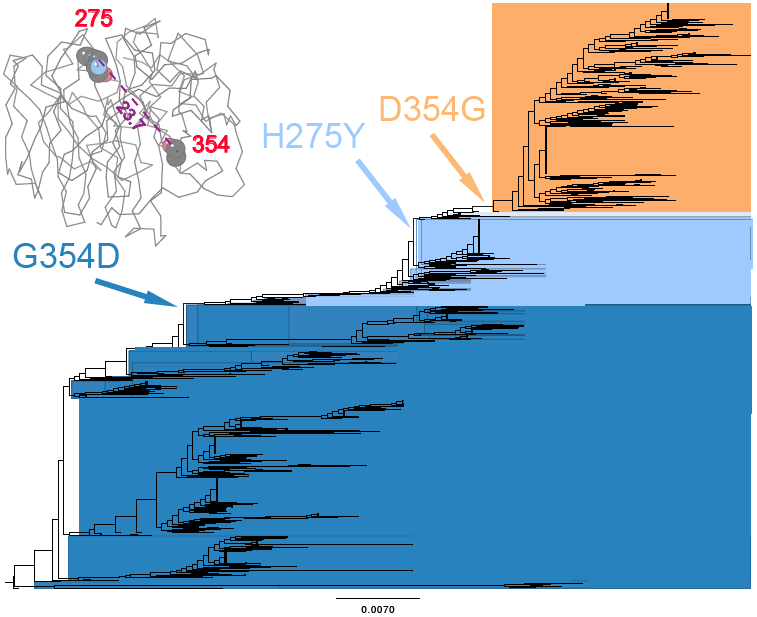
Epistatic pairs of amino acids detected in the Duan14NA data set. The epistatic mutations that we detected are plotted on the NA phylogenetic tree. Inset: the epistatic sites are mapped on three dimensional NA protein structure (based on template 1HA0). The numbers show the AA positions experimentally validated (in red). The numbers in purple show the physical distance between epistatic sites (in Å).

In spite of this negative result, we find that the interacting pair is, again, a long-range interaction (23.7 Å; Figure 3, inset). The objective of the next two sections is to explore more systematically the nature of epistasis in influenza A viruses, focusing more specifically on (i) the prevalence of chained long-range epistasis and (ii) the potential nature of these long-range interactions.

Lastly, we ran our approach on the Li2016 data set. While our analysis found a signal for correlated evolution in 25.7% of the pairs of sites tested by Li *et al*. (2016), 90.0% of our significant pairs of sites were identified as epistatic by Li and collaborators (Figure S19A). We also found a small number of site pairs (37) that were not detected to be epistatic by Li and collaborators. We note however that (i) four of these 37 site pairs could not be statistically tested as the original study had only one biological replicate, (ii) all the site pairs we identify as correlated have epistasis measures that fall within the range of those site pairs deemed significant by Li and collaborators (Figure S19B) and (iii) one site pair we identified forms a Watson-Crick base pair in the folded tRNA molecule. It is therefore possible that the variability in fitness measures in the Li *et al*.’s high throughput experiment be at least partially responsible for these discrepancies. In any case, this comparison supports the results of our simulation studies that show that our method has excellent specificity and mediocre sensitivity.

#### Correlated networks follow a temporal pattern

In a fifth study, the mutations involved in changes of antigenic clusters were investigated, as it was found that double mutations could suffice to explain such cluster changes (Koel *et al*., 2013). The high mutation rate of this virus’ antigen however does not explain why these cluster changes do not occur more often than the observed 3.3 years, which led Koel and collaborators to postulate that “co-mutations” may be required to maintain viral fitness, and hence accelerate evolutionary trajectories when one of the two mutations is neutral or even deleterious (Drake, 2007). In light of our study, the immediate interpretation of these results would be that correlated evolution is involved during such cluster changes. We re-analyzed these data in order to test this hypothesis.

By doing so, we find that some of the substitutions previously (and experimentally) implicated in cluster change are indeed involved in epistasis (Figure 4A-B, Table S10). In particular, as in the Gong13NP data set, we find evidence for networks of correlations, either as short chains such as position 155 interacting with both 158 and 146, which are involved in two consecutive cluster changes, but we also find a much larger network of interactions involving 20 sites, some of which are also involved in the last four cluster changes (Figure 4B). Again, as in the Gong13NP data set, a temporal sequence of substitutions along this network can be found: G124D, found at the SI87/BE89 transition, interacts with K299R, G172D and E82K; G172D interacts with sites involved in the next transition, BE89/BE92, such as G135K, which is again involved in the next transition, BE92/WU95, where G172E interacts with N262S and V196A, which is itself interacting with K156Q, involved in the WU95/SY97 transition; finally, K156Q interacts with T192I, involved in the SY97/FU02 transition. It is tempting to propose that such chained interactions reflect permissive substitutions and hence may provide an explanation, as an evolutionary constraint, to the paradox of high mutation rate and slow antigenic evolution. Yet, can we delve further into the nature of these constraints?

**Figure 4.**
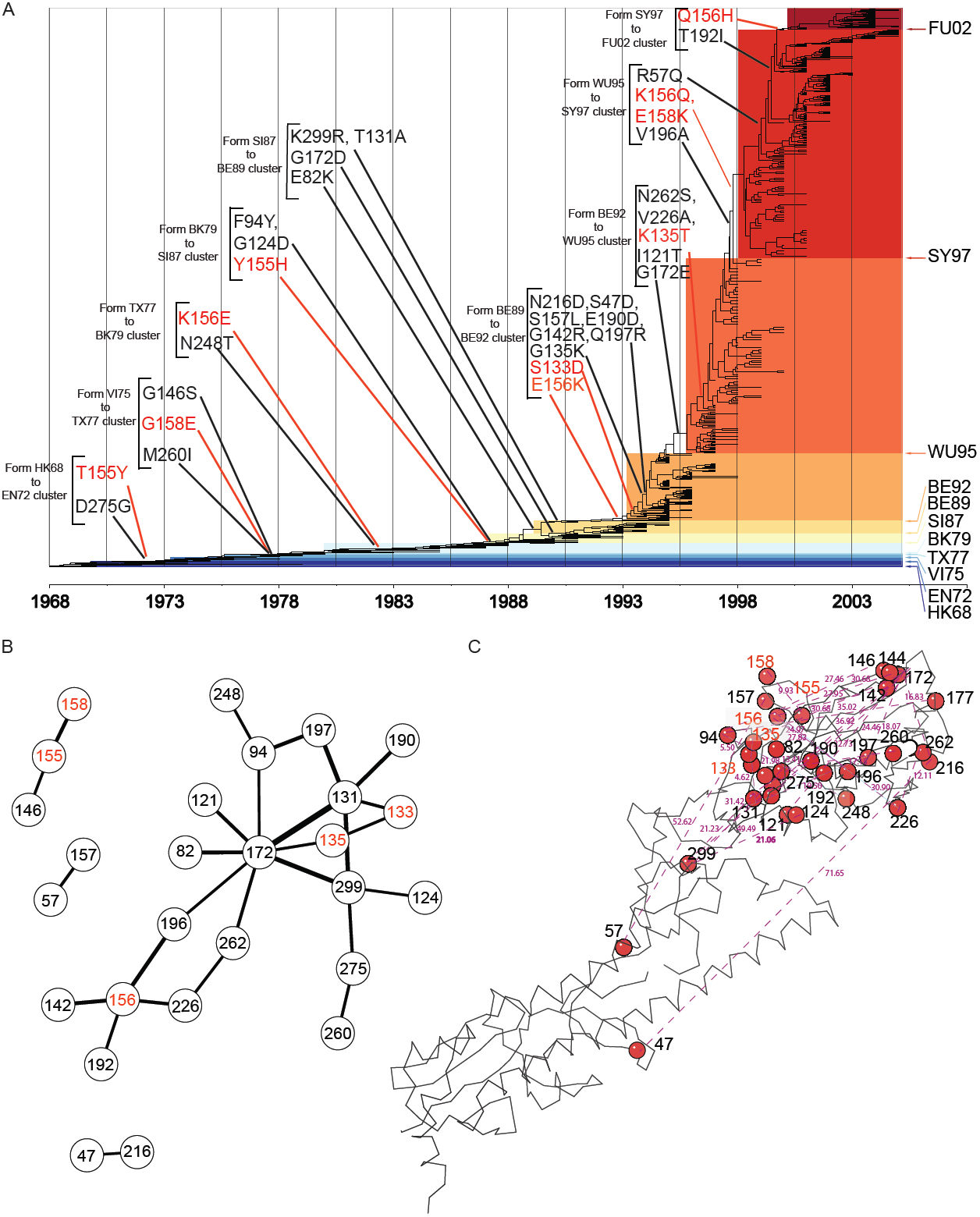
Epistatic pairs of amino acids detected in the Koel13HA data set. (A) The epistatic mutations that we detected are plotted on the HA phylogenetic tree. The substitutions in red were experimentally validated to be responsible for cluster change (Koel *et al.*, 2013). The antigenic clusters are named after the first vaccine strain in the cluster, with letters and digits referring to location and year of isolation (HK, Hong Kong; EN, England; VI, Victoria; TX, Texas; BK, Bangkok; SI, Sichuan; BE, Beijing; WU, Wuhan; SY, Sydney; FU, Fujian). (B) The thickness of the links is proportional to – *log*_10_ *P*-value, the strength of evidence supporting the interaction. (C) The epistatic sites are mapped on three dimensional HA protein structure (based on template 3WHE). The numbers show the AA positions experimentally validated (in red) and those detected only in this study (in black). The numbers in purple show the physical distance between epistatic sites (in Å).

At first inspection, Figure 4C suggests that all these interactions are located in the head of the HA protein and hence might respond to steric constraints, *i.e*., physically-mediated. However, Figure S20 shows that the strength of these associations is not related to physical distance between pairs of interacting sites. Again, the average distance between pairs of interacting sites is 23.7 Å (*sd* = 10.4), which is much larger than the canonical 8.5 Å for close proximity. Can we obtain some evidence about the nature of such long-range interactions?

#### Long-range interactions can be functionally-mediated

The analysis of a sixth data set, Adam03M2, sheds some light on this question. Influenza A viruses are resistant to M2 inhibitors, such as adamantane, and this resistance is associated with the S31N amino acid substitution, which is found in more than 95% of the currently circulating viruses (Wang *et al*., 2013; Garcia and Aris-Brosou, 2014). There is evidence supporting that the spread of S31N may be unrelated to drug selective pressure, but instead results from its interaction with advantageous mutations located elsewhere in the viral genome (Simonsen *et al*., 2007) – or maybe just in the M2 gene. To test this epistatic hypothesis, we used our approach to analyze a data set of M2 sequences.

The results show that only one pair of epistatic sites (S31N×V51I) is detected (Figure 5, Table S11). Again, it is a long-range interaction (35.96 Å), but the reason why this data set is illuminating is that the mutation at position 51 has been shown to play a role in virus replication by stabilizing the amphipathic helixes of the M2 protein (Stewart and Pekosz, 2011). Thus, V51I may enhance the fitness of M2 protein to increase the frequency of adamantane resistance associated with S31N mutation. The reversion I51V that appeared in few sequences in 2000 (red clade in Figure 5) was apparently quickly lost, which supports the hypothesis that V51I mutation is permissive of S31N. This reversion also supports that, even in the face of high mutation rates, (i) our algorithm still maintains high specificity (Figure 1) and (ii) epistasis can be a very powerful force.

**Figure 5.**
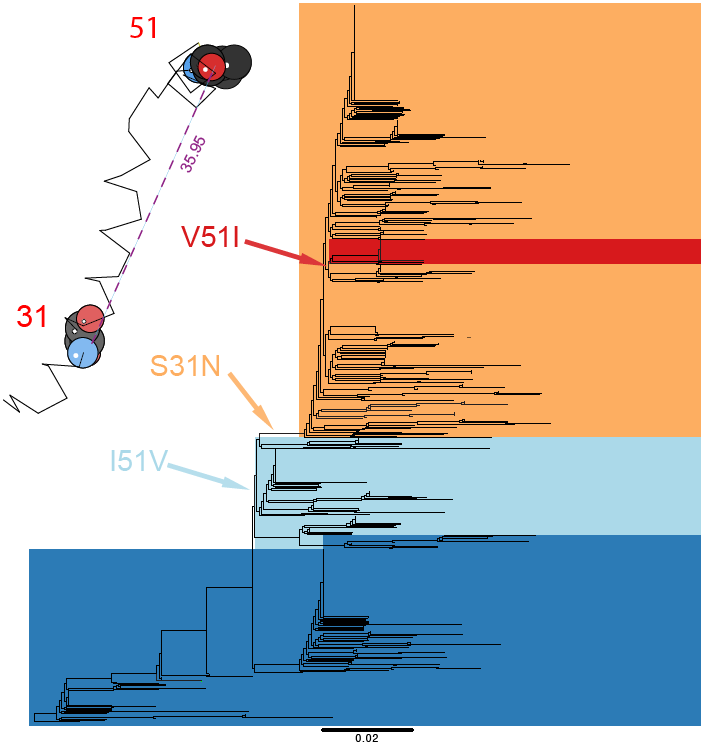
Epistatic pairs of amino acids detected in the Adam03M2 data set. The epistatic mutations that we detected are plotted on the M2 phylogenetic tree. Inset: the epistatic sites are mapped on three dimensional M2 protein structure (based on template 2KIH). The numbers show the AA positions detected (in red). The numbers in purple show the physical distance between epistatic sites (in Å).

## Conclusions

With historical contingency, the accumulation of epistatic substitutions can be seen as a coevolutionary process, where what happens at one AA site is contitional on what happened at another site. Here, it is this very idea of coevolution that we harnessed by co-opting a method developed in ecology to test for the correlated evolution of phenotypic traits (Pagel, 1994; Pagel and Meade, 2006): instead of treating pairs of phenotypic traits as such, we repurposed the method to deal with pairs of AA sites. Because the original method was developed to handle binary traits, we explored two ways of recoding data and showed that treating AAs as outgroup/ingroup consensus states was a more sensible (albeit less intuitive) option than using physicochemical properties. Extensive simulations demonstrated mediocre sensitivity, but excellent specificity and precision, even in the face of dinucleotide bias, so that pairs of AAs that are detected can be assumed to be actually coevolving. We then validated our approach against other computational results, showing marginal overlap, and against experimentally-validated results, showing extensive overlap, hereby suggesting that detected pairs of AAs are genuinely interacting.

With this good statistical behavior, further analyses of independent influenza data sets showed a consistent pattern: (i) many pairs of correlated sites are involved in epistatic interactions, (ii) these pairs of sites form extensive networks of sites, consistent with Poon *et al*. (2007b), but that are also affected by substitutions that occur sequentially – showing evidence for historical contingency in fast-evolving organisms -and (iii), more intriguingly, that these epistatic pairs of sites form long-range spatial interactions. This latter point precludes the idea of a close physical link as in the case of tRNA molecules (Kimura, 1985; Chen *et al*., 1999), so that these long-range interactions must bring about stability (Thomas *et al*., 2010) and/or conformational (Mitraki *et al*., 1991; Newcomb *et al*., 1997; Harms and Thornton, 2014) and/or functional changes, as in the case of the M2 data set or in the case of the Ebola virus (Ibeh *et al*., 2016). While there is evidence that epistasis can be prevalent in RNA viruses (at least 31% in (Shapiro *et al*., 2006)) and in bacteria (15% in (Weinreich *et al*., 2006)), it is not impossible that epistasis reflects an evolutionary constraint stronger in RNA viruses than in organisms with larger and more redundant genomes: because these viruses have a small genome, mutations are expected to have large fitness effects, which can be alleviated by compensatory mutations (Sanjuán and Elena, 2006).

Our choice of focusing on a segmented RNA virus such as influenza may be problematic, in particular in the case of H3N2 viruses, which show a pattern of punctuated evolution that can be interpreted as the result of clonal interference (Illingworth and Mustonen, 2012; Strelkowa and Lässig, 2012). In the absence of recombination, each segment of the virus evolves as a clone, and each clone accumulates different beneficial mutations, only one of which becomes fixed, alongside hitchhiking deleterious mutations that, in our context, would show a pattern of correlated evolution. While (i) this process has to date only be found in H3N2 viruses, and (ii) we also find evidence for chained epistasis in H1N1 viruses (KDBP11-H1, KDBP11-N1, Duan14NA) as well as in eukaryotic tRNA gene (Li2016), clonal interference remains a problem for our approach. On the other hand, the use of influenza has allowed us to limit our analysis to searching for evidence of epistasis within genes, *contra* among genes. This way of analyzing data, intra-genically, might make sense when different segments code for proteins involved in relatively different functions. However, many RNA viruses have overlapping reading frames (such as the M and NS segments in influenza A), so that we can also expect coevolution across viral genes (Neverov *et al*., 2015). Furthermore, experimental evidence in other organisms, such as yeasts, shows that epistatic interactions can involve multiple genes and hence be *inter*-genic (Sorrells *et al*., 2015). Although a method to detect epistasis in segmented genomes was proposed (Neverov *et al*., 2015), the computational costs of whole-genome scans can seem prohibitive. Assessing the prevalence of epistasis over entire genomes remains an unexplored area − but one that we are currently investigating.

In this context, how can we explain the existence of large networks of interacting sites? One possibility would be that a changing environment creates new adaptive landscapes, and that natural populations (*contra* those from *in vitro* studies) do not climb peaks on the landscape but rather chase moving targets (Gavrilets, 2004, p. 36). While it is not clear whether such landscapes are robust to changing environments (Hartl, 2014), they are certainly a reality in the world of viruses, where vaccination regularly alters the adaptive landscape, hereby leading to chained networks of epistatic interactions – as a mere by-product of evolution. But then, one can wonder if the metaphor of a landscape itself is appropriate when all mutational trajectories are not accessible from specific genomic backgrounds (Weinreich, 2010; Sorrells *et al*., 2015). This may be one of the reasons why evolution is so difficult to predict (Weinreich, 2010; Sandie and Aris-Brosou, 2014; Natarajan *et al*., 2016).

## Acknowledgments

We thank Jeremy Dettman, Neke Ibeh, Rees Kassen, Nicolas Rodrigue and Alex Wong for fruitful discussions, Sergey Kryazhimskiy and Joshua Plotkin as well as Chuan Li and Jianzhi Zhang for sharing data with us, and two anonymous reviewers for comments that improved the article. We are grateful to Compute Canada and to Ontario’s High Performance Computing Virtual Laboratory for giving us compute time. This work was supported by the Natural Sciences Research Council of Canada, the Canada Foundation for Innovation (SAB), and by the University of Ottawa (JCN, JD).

